# Integrated analysis of intestinal microbiota and metabolomic reveals that decapod iridescent virus 1 (DIV1) infection induces secondary bacterial infection and metabolic reprogramming in *Marsupenaeus japonicus*

**DOI:** 10.1101/2022.06.27.497879

**Authors:** Zihao He, Yunqi Zhong, Minze Liao, Linxin Dai, Yue Wang, Shuang Zhang, Chengbo Sun

## Abstract

In recent years, with global warming and increasing marine pollution, some novel marine viruses have become widespread in the aquaculture industry, causing huge losses to the aquaculture industry. Decapod iridescent virus 1 (DIV1) is one of the newly discovered marine viruses that has been reported to be detected in a variety of farmed crustacean and wild populations. Previous studies have found that DIV1 can induce the Warburg effect. To further explore the effect of DIV1-induced metabolic reprogramming on *Marsupenaeus japonicus* intestinal metabolome and microbiota and the consequence on immune response, histological analysis, enzyme activity analysis and the integrated analysis of intestinal microbiome and metabolomics were performed in this study. The results showed that obvious injury in the intestinal mucosa was observed after DIV1 infection. The oxidative and antioxidant capacity of the shrimp intestine was unbalanced, the activity of lysozyme was decreased, and the activities of digestive enzymes were disordered, causing secondary bacterial infection. In addition, the increased abundance of harmful bacteria, such as *Photobacterium* and *Vibrio*, synergized with DIV1 to promote the Warburg effect and induce metabolic reprogramming, thereby providing material and energy for DIV1 replication. This study is the first to report the changes of intestinal microbiota and metabolites of *M. japonicus* under DIV1 infection, demonstrating that DIV1 can induce secondary bacterial infection and metabolic reprogramming, and several highly related bacteria and metabolites were screened as biomarkers. These biomarkers can be leveraged for diagnosis of pathogenic infections or incorporated as exogenous metabolites to enhance immune response.

## 1. Introduction

Viruses are an important part of marine ecosystems. With the development of genetic technology, more and more new marine RNA and DNA viruses have been discovered (Gregory et al., 2019; Zayed et al., 2022). Despite their tiny size, viruses play a large role in marine ecosystems and food webs. On the one hand, marine viruses can infect a variety of oceans organisms, lysing their cells and releasing carbon and other nutrients that impact the food web (Suttle et al., 2007). On the other hand, marine viruses often contain host-derived metabolic genes (i.e., auxiliary metabolic genes; AMGs), which are hypothesized to increase viral replication and alter ecosystem-level productivity through reprogramming host metabolism (Hurwitz and U’Ren., 2016). To date, several crustacean viruses have been found to induce metabolic reprogramming to promote survival and replication, including white spot syndrome virus (WSSV) (Chen et al., 2011; Hsieh et al., 2015; Apún-Molina et al., 2017; Godoy-Lugo et al., 2019), Taura syndrome virus (TSV) (Zeng et al., 2014) and IHHNV (Galvan-Alvarez et al., 2012). In recent years, with global warming and increasing marine pollution, some novel marine viruses have become widespread in the aquaculture industry, causing huge losses to the aquaculture industry (He et al., 2022). Decapod iridescent virus 1 (DIV1) is one of the newly discovered marine viruses. Since China initiated the DIV1 surveillance in 2017, DIV1 has been detected in shrimp culture ponds in several provinces (Liao et al., 2022). In addition to farmed shrimp, in 2020, Srisala et al. also detected DIV1 in wild populations of *Penaeus monodon* in the Indian Ocean (Srisala et al., 2021). Up to now, DIV1 has been known as a highly lethal virus with global risk of transmission, capable of infecting freshwater and marine crustaceans, including *Marsupenaeus japonicus* (He et al., 2021a).

Kuruma shrimp *M. japonicus* is widely distributed in the Indo-Western Pacific region and the East and South China seas (Dall et al., 1990). Due to its high economic value, strong environmental adaptability and suitability for long-distance transportation, it has now become one of the most farmed prawns in China (He et al., 2020; Yuan et al., 2016). Our previous study demonstrated that *M. japonicus* was a susceptible host to DIV1 (He et al., 2021a; He et al., 2022). Through mRNA-seq and miRNA-seq analyses and their association analysis, we preliminarily revealed the hemocyte and intestinal immune response of *M. japonicus* to DIV1 infection (He et al., 2021a; He et al., 2022), and proposed that DIV1 can promote the Warburg effect by regulating host miRNA and mRNA expression. The Warburg effect was also observed in DIV1-infected *Litopenaeus vannamei* and *Penaeus monodon*. However, only the miRNA and mRNA espression profiles under DIV1 infection were revealed, the deep mechanism of the immune response needs further analysis, especially the intestinal immunity.

As an important immune and digestive organ of shrimp, the intestine and symbiotic microorganisms together constitute a complex ecosystem, which plays an important role in maintaining the function of the shrimp immune system. The intestinal immune system of shrimp is mainly composed of three parts, including the mechanical barrier formed by the tight junction and adhesion junction of intestinal mucosal cells, the innate immune barrier formed by intestinal hemocytes and their secreted immune factors, and the biological barrier formed by the intestinal microbiota and their secretions (Zheng et al., 2016). The potential cooperation of bacteria and viruses in promoting disease development has received extensive attention from researchers in recent years. Numerous studies have shown that viral infection can alter the composition and function of the shrimp intestinal microbiota, induce secondary bacterial infection, and impair host immunity (Wang et al., 2018; Chen et al., 2021; Niu et al., 2022). However, similar to other invertebrates, shrimp lack specific immunity and cannot acquire antibodies through vaccination. Therefore, regulating host metabolic reprogramming by exogenous metabolites to enhance host immunity and ability to withstand environmental has become one of the best strategies for shrimp to resist viral infection (Wang et al., 2018; Wu et al., 2021; Wang et al., 2021). It will be interesting to explore the effect of metabolic reprogramming on host intestinal metabolome and microbiota and the consequence on immune response. To date, there are no reports on the effects of DIV1 infection on the intestinal metabolome and microbiome of shrimp.

To gain a more comprehensive understanding of the interaction between DIV1 and shrimp, this study is the first to investigate the changes of intestinal microbiota and metabolites of *M. japonicus* under DIV1 infection through the integrated analysis of intestinal microbiome and metabolomics, demonstrating that DIV1 can induce secondary bacterial infection and metabolic reprogramming, and several highly related bacteria and metabolites were screened as biomarkers. The results are beneficial to provide a theoretical basis for virus control technology.

## 2. Materials and Methods

### 2.1. Shrimp and Rearing Conditions

Healthy *M. japonicus* with an average body weight of 10.5 ± 1.6 g were randomly obatained from a local farming pond in East Island Marine Biological Research Base, Guangdong Ocean University (Zhanjiang, China). The shrimp were randomly sampled and tested by PCR to ensure that they were free from WSSV, IHHNV, and DIV1. The detection method was consistent with the previous study (He et al., 2021b). Every 30 *M. japonicus* were acclimatized in 0.3-m^3^ tanks with aerated and filtered seawater (salinity 30‰, pH 7.5, temperature 28°C) for 7 days before the DIV1 challenge experiment. Commercial feed was provided to the shrimp daily at 5% of their body weight three times per day, and nearly 90% of the seawater was exchanged per day.

### 2.2. DIV1 Challenge and Sample Collection

After 7 days of acclimation, the healthy shrimp were divided into two groups: the control group and the DIV1-infected group. Each group included three replicate tanks, and each tank containing 30 individuals. Base on LC_50_ test results from previous studies (He et al., 2021a), we set the DIV1 injection concentration of this study as 3.95 × 10^9^ copies/μg DNA. Each *M. japonicus* from the DIV1-infected group was intramuscularly injected with 50 μL of DIV1 inoculum, while each M. japonicus from the negative control group was intramuscularly injected with 50 μL of phosphate-buffered saline (PBS; pH 7.4). The methods of viral inoculum preparation and quantification can be found in previous studies (Liao et al., 2020).

Based on previous research (He et al., 2021a), the *M. japonicus* at twenty-four hour post-injection (hpi) were collected as samples under an aseptic condition. The intestines from 3 random individuals in the same group were combined as one sample and immediately frozen in liquid nitrogen before storing at -80°C until experimental analysis. In detail, each group had 6 samples were used for enzyme activity analysis, 6 samples were used for intestine microbiome analysis and 6 samples were used for intestinal metabolomics analysis. For histological analysis, the intestines of 6 shrimp per group were used to assess intestine tissue damage. Intestines of these shrimp were fixed in 10% formalin.

### 2.3. Histological Analysis

After fixed in 10% formalin at 4°C for 24 h, the fixed intestine were dehydrated in gradient ethanol, hyalinized in xylene, and embedded in paraffin wax. Next, the paraffin blocks were sectioned at 5-μm thickness. The sections were colleted on glass slides and stained with hematoxylin and eosin (H&E). The intestinal sections were examined by a microscope.

### 2.4. Enzyme Activity Analysis

Enzymatic biomarkers of functional responses in the intestine were measured using commercial detection kits (Jianglai Bioengineering Institute, Shanghai, China) according to protocols of the manufacturer. Prior to analysis, 6 samples in each group were were homogenized in pre-chilled PBS (1:9 dilution) and then centrifuged for 10 min (4°C and 5,000 × g) to obtain the supernatant for further use. Superoxide dismutase (SOD), catalase (CAT) and lysozyme (LYZ) activities were used to reflect the nonspecific immunity in the shrimp intestine. α-Amylase (α-AMS), lipase (LPS) and trypsin (TPS) activities were used to reflect the digestive function of shrimp.

### 2.5. Intestinal Microbiome Analysis

Total bacterial DNA was extracted by the *EasyPure*® Marine Animal Genomic DNA Kit (TransGen Biotech, China) following the manufacturer’s directions. The concentration and purity of total DNA were determined by SimpliNano (GE Healthcare, United States) and 1% agarose gels. The primers pair 515F (5′-GTGCCAGCMGCCGCGG-3′) and 806R (5-′GGACTACNNGGGTATCTAAT-3′) were used to amplify the V4 hypervariable region of 16S rRNA gene, which was modified with a barcode tag with a random 6-base oligos. PCR amplification reaction system and parameters can refer to previous studies (Xiong et al., 2012). After the PCR products were purified and mixed in equidensity ratios, sequencing libraries were constructed, and then sequenced by BGI (Shenzhen, China) with the Illumina Genome Analyzer technology. All raw sequencing data of intestinal microbiota was submitted to the Sequence Read Archive (SRA) (accession: PRJNA720257).

Sequences from raw data were analyzed and filtered by QIIME (v1.8.0). Sequence analysis was performed with UPARSE software (v7.0.1090), and the operational taxonomic units (OTUs) were defined with ≥ 97% similarity. Chimeric sequences were identified with UCHIME (v4.2.40). Alpha diversity was calculated using mothur software with five metrics, including the observed OTUs, Chao1, ACE, Shannon and Simpson indexes. Rarefaction curves were generated based on these metrics. The shared and unique OTUs between two groups were figured out by a Venn diagram. Beta diversity index based on the phylogenetic relationship between OTUs was used to calculate the Unifrac distance (unweighted Unifrac) and the results of Beta diversity were showed through PCoA and UPGMA Phylogenetic Dendrogram. A bar plot of the microbial community was constructed at the phylum, family and genus level respectively. To study the functional characteristics of bacterial communities, Kyoto Encyclopedia of Genes and Genomes (KEGG) functions were predicted using the PICRUSt software.

### 2.6. Intestinal Metabolomics Analysis

Six intestinal samples replicates of shrimp from each group were used for metabolomic analysis. All the samples were taken from the refrigerator at -80°C and whawed in the refrigerator at 4°C, and metabolite extraction was performed using methanol and 2-chlorobenzalanine. Twenty microlitres of each sample was taken for quality control (QC), and the rest was used for LC-MS detection. Liquid chromatography was accomplished in a Thermo Ultimate 3000 system equipped with an ACQUITY UPLC® BEH C18 (1.7 μm 2.1 × 100 mm, Waters, USA) column. Mass spectrometry was executed on a Q Exactive HF mass spectrometer (Thermo Fisher Scientific, USA). Data-dependent acquisition (DDA) MS/ MS experiments were performed with HCD scans. Dynamic exclusion was implemented to remove some unnecessary information in the MS/MS spectra.

Peak-identification, peak-alignment and compound identification was conducted using Compound Discoverer software (v3.1). All the data were determined using quality control (QC) and quality assurance (QA). Partial least squares-discriminant analysis (PLS-DA) of the metabolomics data was performed using the R language ropls package. All the metabolites were classified according to Kyoto Encyclopedia of Genes and Genomes (KEGG) and Human Metabolome Database (HMDB). The PLS-DA model was used to determine the differential metabolites (DMs) between the control group and the DIV1-infected group with the first principal component of variable importance in projection (VIP) values (VIP ≥ 1) combined with a *q*-value ≤0.05. The DMs were annotated with KEGG pathway analysis to further identified the change characteristics of the functional metabolites related to the immunity of the shrimp.

### 2.7. Correlation Analysis of Intestinal Bacteria and DMs

Canonical correlation analysis and spearman correlation analysis were employed to reveal the correlation between significantly altered intestinal bacteria and intestinal DMs, and the results were shown with scatterplots and heat maps. The thresholds for correlation coefficients and *p*-values were not set. *p* < 0.05 was regarded as statistically significant, *p* < 0.01 was regarded as very significant, and *p* < 0.001 was regarded as extremely significant.

## 3. Result

### 3.1. Intestinal Histological Changes

Histological analysis of the intestinal sections from the control group and the DIV1-infected group are shown in Figure 1. The H&E staining showed that no histological changes were observed in the controls (Figure 1A). Contrastively, the microstructure of the intestine in the DIV1-infected group had lesions, and the intestinal epithelial cells were completely detached from the basement membrane and severely destroyed (Figure 1B).

**Figure 1.**
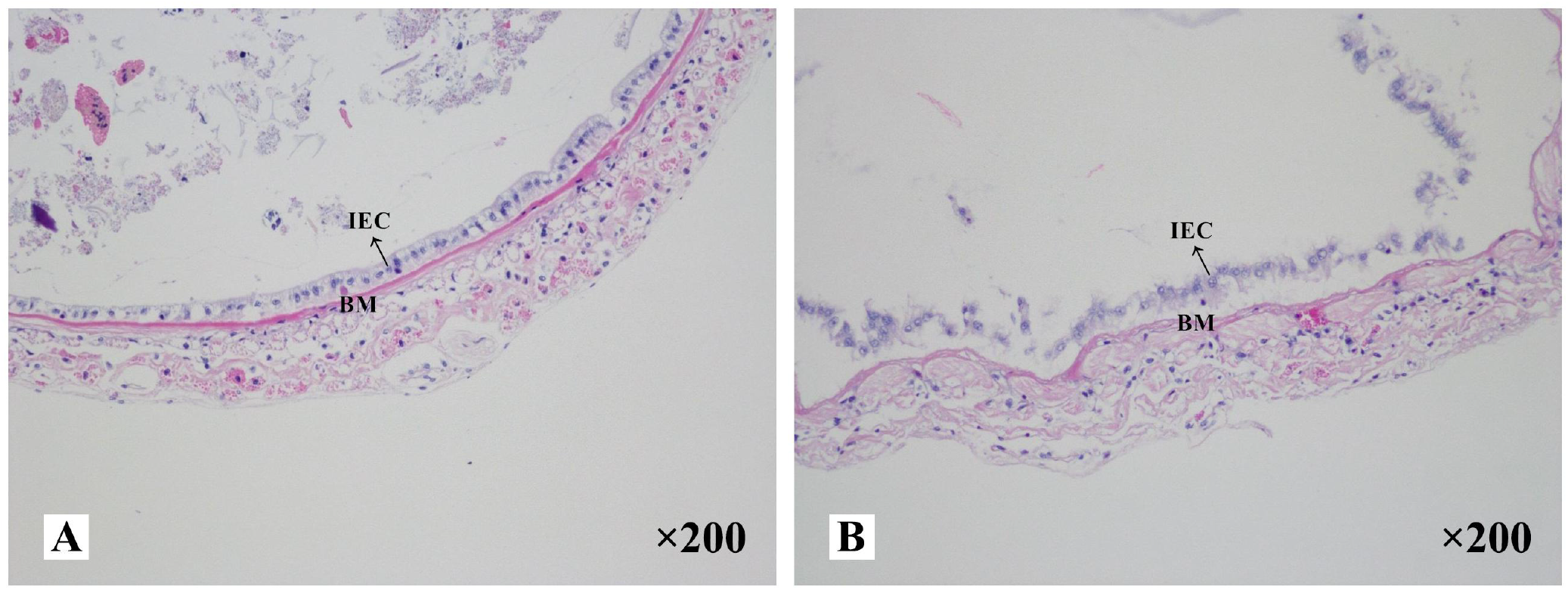
Photomicrographs of intestinal sections in (A) the control group and (B) the DIV1-infected group. The magnification was ×200. IEC: intestinal epithelial cells; BM: basement membrane.

### 3.2. Immune and Digestive Enzymes Activity Abnormalities

At 24 hpi, the activities of SOD, CAT and LYZ in the intestine were detected to evaluate the effect of DIV1 infection on the nonspecific immunity of *M. japonicus* (Figure 2A,B&C), and the activities of α-AMS, LPS and TPS in the intestine were detected to evaluate the digestive function (Figure 2D,E&F). Compared with the control group, the activity of SOD, α-AMS and LPS were significantly increased (*p* < 0.001), while the activities of CAT, LYZ and TPS were significantly decreased (*p* < 0.001). It was worth noting that, after infection with DIV1, both the immune and digestive enzymes in the intestine were extremely significantly altered (*p* < 0.001).

**Figure 2.**
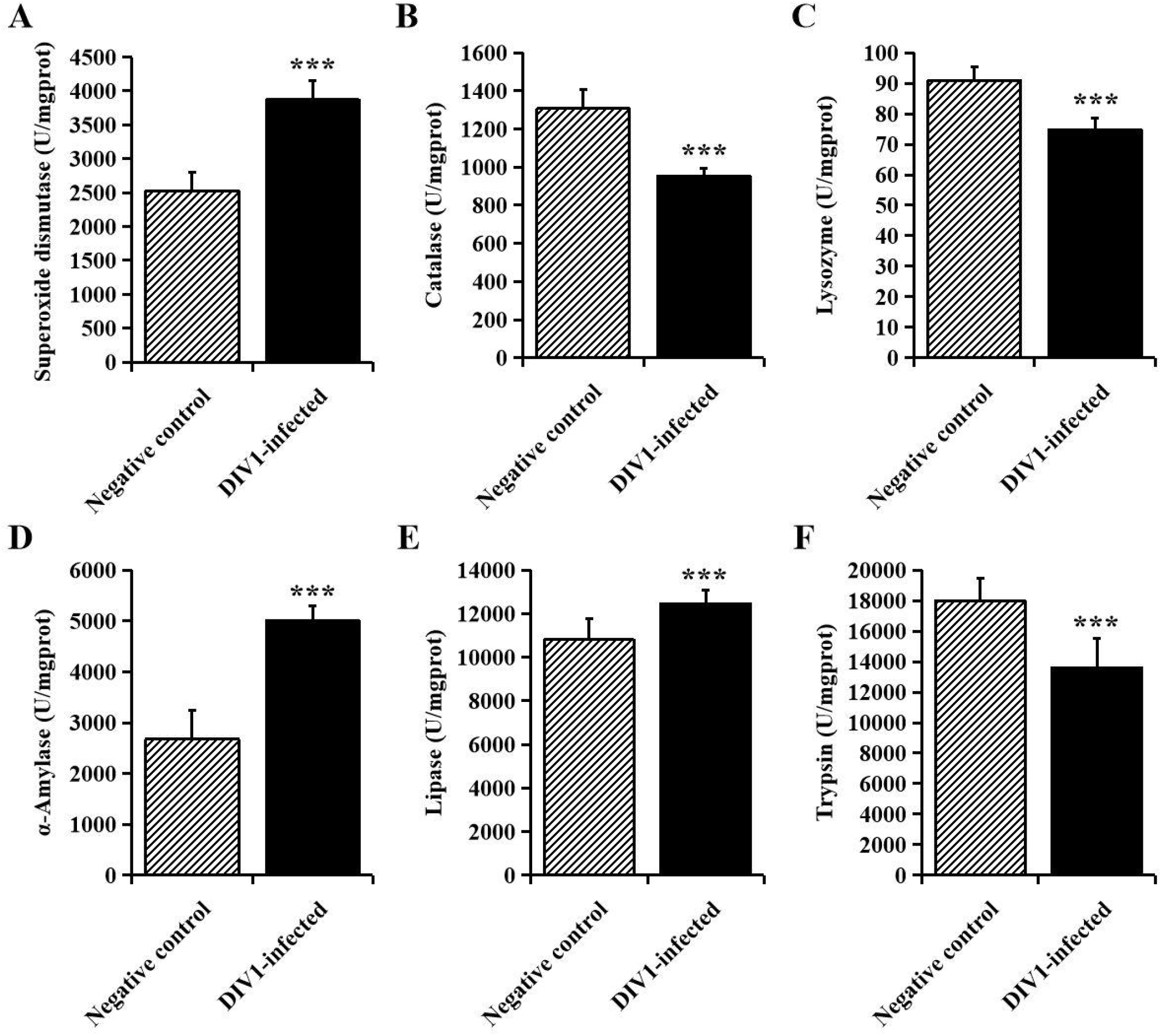
The activities of immune and digestive enzymes in the intestine of *M. japonicus* (mean ± SD). **(A)** Superoxide dismutase (SOD). **(B)** catalase (CAT). **(C)** lysozyme (LYZ). **(D)** α-Amylase. **(E)** lipase (LPS) and **(F)** trypsin (TPS). The statistically significant differences between the two groups were calculated by Student’s *t*-test (\**p* < 0.05, \*\**p* < 0.01, \*\*\**p* < 0.001).

### 3.3. Intestinal Microbiota Changes

#### 3.3.1. Richness and Diversity

A total of 711,264 high-quality sequences were generated from 12 intestinal samples. The clean reads ranged from 69,929 to 63,969, with an average of 59,272 clean reads per sample. After the alignment, the sequences of the control group and the DIV1-infected group were clustered into 1,053 and 1,057 OTUs respectively with a 97% sequence similarity. Among them, there were 421 unique OTUs in the control group, 425 unique OTUs in the DIV1-infected group, and 632 OTUs were the same in both the control group and the DIV1-infected group (Figure 3A). A rarefaction curve analysis of the observed species per sample was sufficient (Figure 3B). To investigate the differences of species diversity and richness between two groups, the alpha diversity indexes were calculated, including observed OTUs, Chao1, ACE, Shannon and Simpson indexes, ranging from 270 to 715, 400.25 to 753.01, 424.50 to 765.22, 0.41 to 3.19 and 0.10 to 0.89, respectively (Table 1). Community richness indexes (observed OTUs, Chao1, ACE) were not significantly changed, while community diversity indexes (Shannon and Simpson) were significantly changed (*p* < 0.05). Beta diversity analysis was performed to comparative analyze the similarity and difference of intestinal microbial community in different groups. The PCoA with unweighted unifrac distance was further performed to confirm that intestinal bacteria in the control group and the DIV1-infected group were clearly separated and samples from the same group were clustered closer (Figure 3C). UPGMA Phylogenetic Dendrogram showed that all detected samples were divided into two main clades, and the intestinal microbial in the same group had a high degree of similarity (Figure 3D).

**Figure 3.**
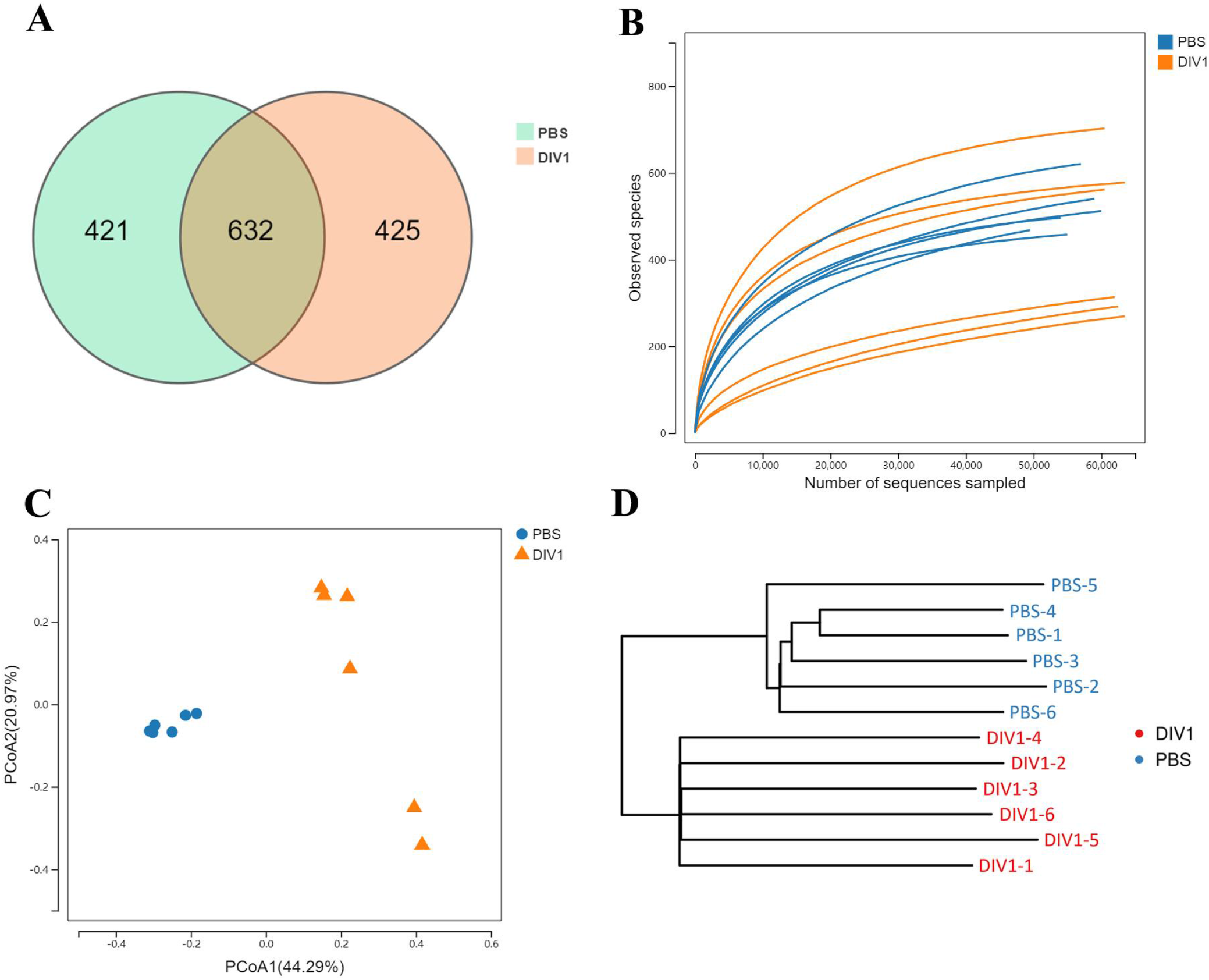
Intestinal microbial richness and diversity of *M. japonicus* after DIV1 infection. **(A)** Veen diagram showing the unique and shared OTUs of intestinal microbiota in the control groups and the DIV1-infected group. **(B)** Rarefaction curves of OTUs clustered at 97% sequence identity different samples. **(C)** PCoA plot and **(D)** UPGMA Phylogenetic Dendrogram with unweighted unifrac distance showed the microbial diversity of samples.

**Table 1.** Alpha diversity index analysis in the control group and the DIV1-infected group.

| Sample | Sobs | Chao1 | ACE | Shannon | Simpson | Coverage |
| --- | --- | --- | --- | --- | --- | --- |
| PBS-1 | 512 | 574.06 | 577.24 | 2.65 | 0.18 | 0.998304 |
| PBS-2 | 497 | 529.76 | 538.83 | 2.06 | 0.37 | 0.998602 |
| PBS-3 | 458 | 490.50 | 493.66 | 2.40 | 0.30 | 0.998809 |
| PBS-4 | 541 | 640.29 | 666.17 | 2.60 | 0.21 | 0.997645 |
| PBS-5 | 469 | 580.91 | 605.59 | 2.17 | 0.22 | 0.997116 |
| PBS-6 | 621 | 678.58 | 703.00 | 3.18 | 0.10 | 0.997910 |
| DIV1-1 | 703 | 736.11 | 747.55 | 2.73 | 0.27 | 0.998572 |
| DIV1-2 | 292 | 470.22 | 589.70 | 0.37 | 0.90 | 0.997863 |
| DIV1-3 | 314 | 450.56 | 537.92 | 0.74 | 0.80 | 0.998154 |
| DIV1-4 | 562 | 643.01 | 639.31 | 2.10 | 0.34 | 0.998189 |
| DIV1-5 | 270 | 404.72 | 430.81 | 0.76 | 0.64 | 0.998124 |
| DIV1-6 | 578 | 611.13 | 614.32 | 2.04 | 0.43 | 0.998890 |
| *p*-values | 0.441 | 0.631 | 0.940 | 0.032 | 0.014 | 0.451 |

#### 3.3.2. Intestinal Microbial Composition

The taxa of dominant bacteria among the two groups were similar, while their abundance was altered significantly. At the phylum level, compared with the control group, the relative abundance of Proteobacteria was significantly increased in the DIV1-infected group (*p* < 0.01), while the relative abundance of Actinobacteria and Cyanobacteria were significantly decreased (*p* < 0.01) (Figure 4A). At the family level, the relative abundance of Vibrionaceae and Sapropiraceae were significantly increased in the DIV1-infected group (*p* < 0.05), while the relative abundance of Corynebacteriaceae, Rhodobacteraceae and Enterobacteriaceae were significantly decreased (*p* < 0.05) (Figure 4B). At the genus level, differences were also observed. The relative abundance of *Photobacterium* was significantly decreased (*p* < 0.01), while the relative abundance of *Corynebacterium, Sagittula* and *Morganella* were significantly decreased (*p* < 0.05).

**Figure 4.**
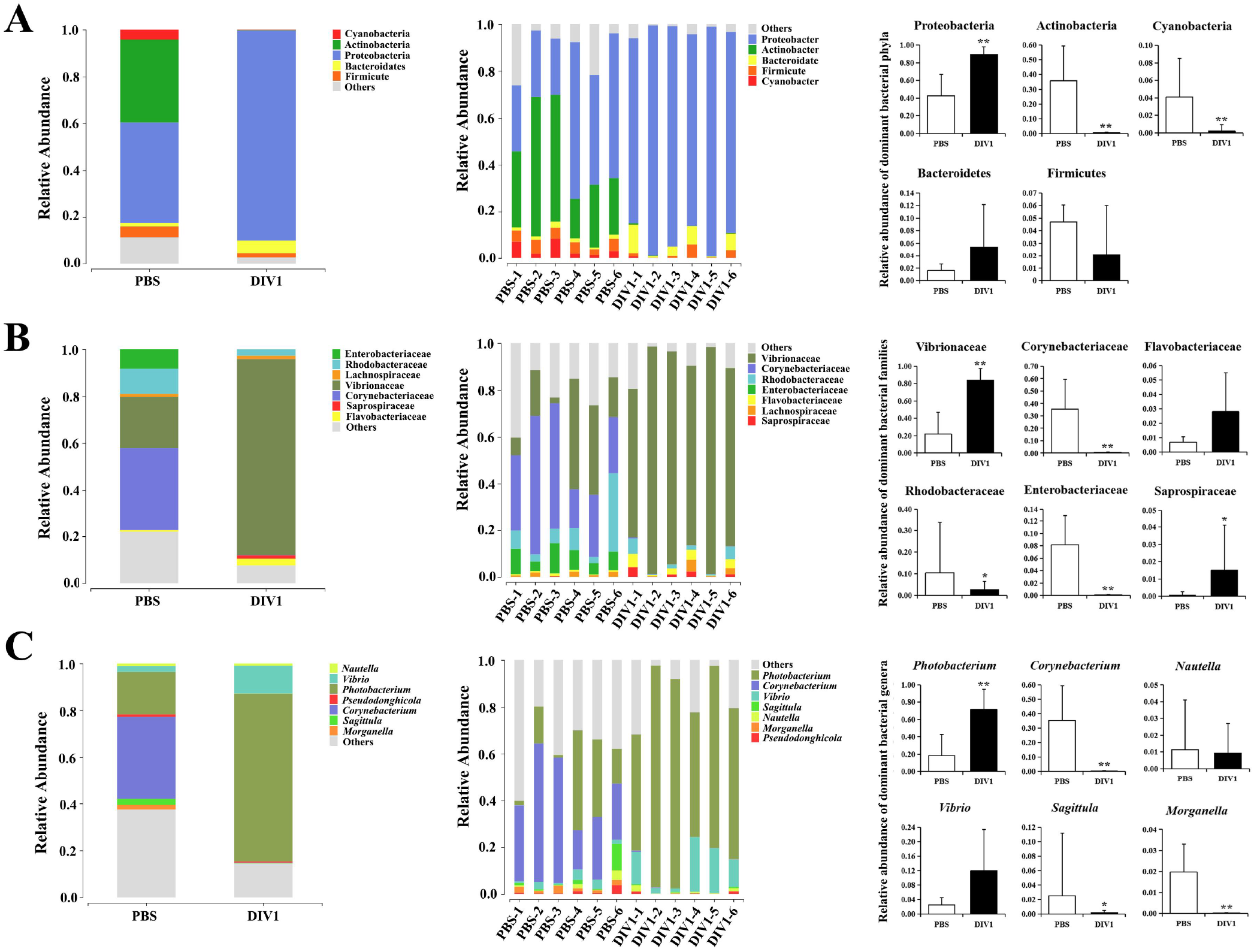
Structure and composition of the intestinal bacterial communities under DIV1 infection (mean ± SD). **(A)** Relative abundances of dominant bacterial phyla. **(B)** Relative abundances of dominant bacterial families. **(C)** Relative abundances of dominant bacterial genera.

#### 3.3.3. Changes in the intestinal bacterial phylotypes

LEfSe was employed to analyse the differential abundances of bacterial taxa in the two groups. In the cladogram, the families Vibrionaceae and Saprospiraceae were enriched in the DIV1-infected group, and Corynebacteriaceae, Rhodobacteraceae, Enterobacteriaceae and Lactobacillaceae were enriched in the control group (Figure 5A). With an LDA score greater than 4.0, *Photobacterium* and *Vibrio* dominated in the DIV1-infected group, and Corynebacterium and Ruegeria dominated in the control group (Figure 5B). The prediction function of the intestinal microbiota was analysed using PICRUSt. Result showed that the relative abundance of “Carbohydrate metabolism”, “Metabolism of cofactors and vitamins” and “Amino acid metabolism” were the top 3 in the two groups. It was worth noting that the relative abundance of “infectious diseases: bacterial” was significantly increased under DIV1 infection (*p* < 0.01).

**Figure 5.**
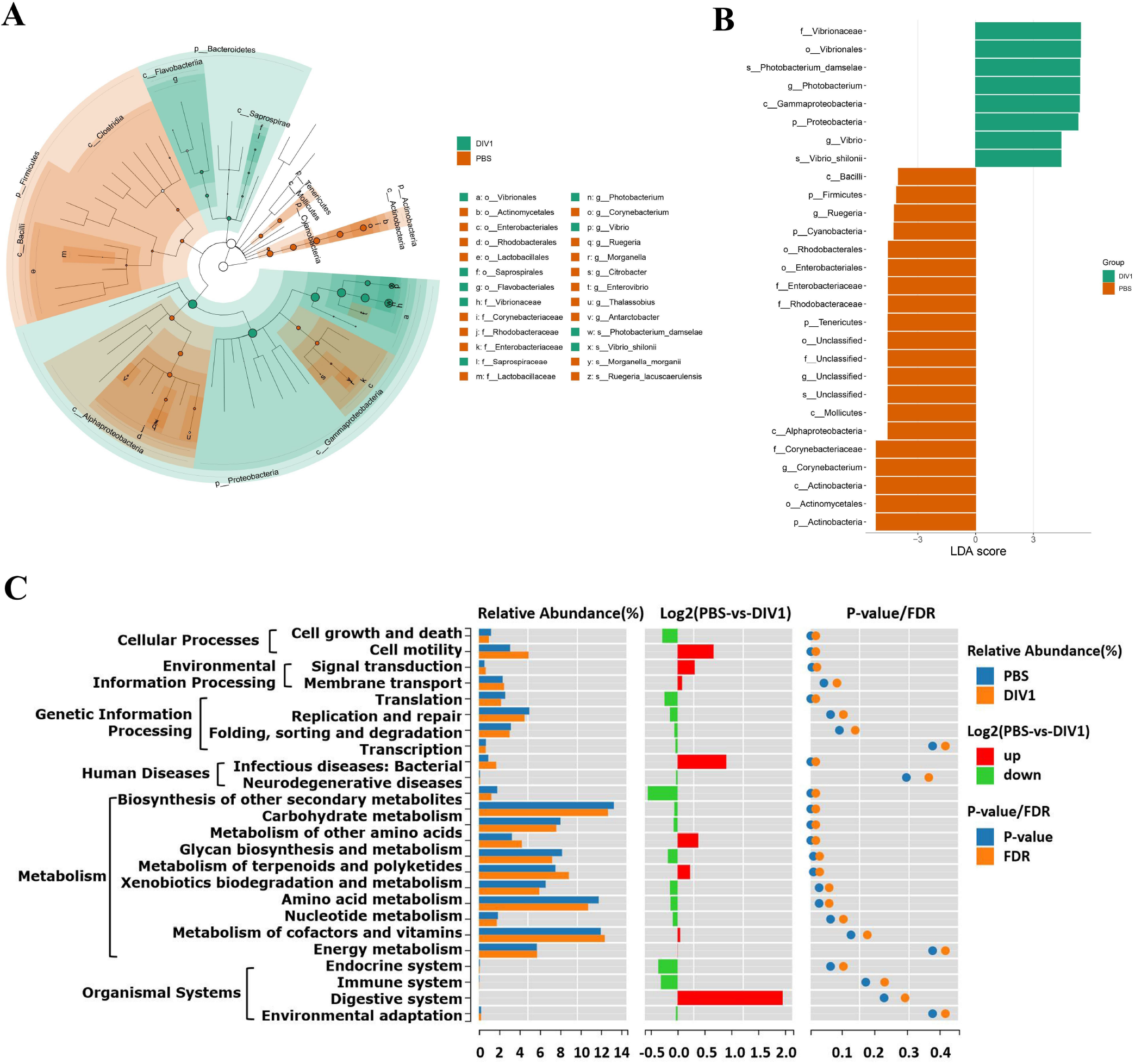
Intergroup variation, functional analysis of intestinal microbiota of *M. japonicus* after DIV1 infection. **(A)** LEfSe cladogram. **(B)** LDA score of LEfSe-PICRUSt. **(C)** Microbial metabolism prediction based on KEGG pathway analysis.

### 3.4. Intestinal Metabolic Pattern Alterations

#### 3.4.1. Multivariate analysis of the metabolite profiles

Metabolomic analysis was conducted to explore the alterations in intestinal metabolic profiles after DIV1 infection. The PLS-DA score plot and permutation test showed a significant different between the two groups in both positive and negative ionization modes (Figure S1), suggesting that DIV1 infection caused metabolic phenotype alterations in shrimp intestine. A total of 3,009 metabolites were identified in the shrimp intestine (including 2,096 metabolites were identified in the positive ion mode and 913 metabolites were identified in the negative ion mode). The classification results of the identified metabolites were shown in Figure 6. The largest category in the positive ion mode was “amino acids, peptides, and analogues” (253 metabolites), followed by “fatty acyls” (147 metabolites) and “benzene and derivatives” (139 metabolites) (Figure 6A), and the largest category in the negative ion mode was “amino acids, peptides, and analogues” (132 metabolites), followed by “fatty acyls” (58 metabolites) and “organic acids” (29 metabolites) (Figure 6B).

**Figure 6.**
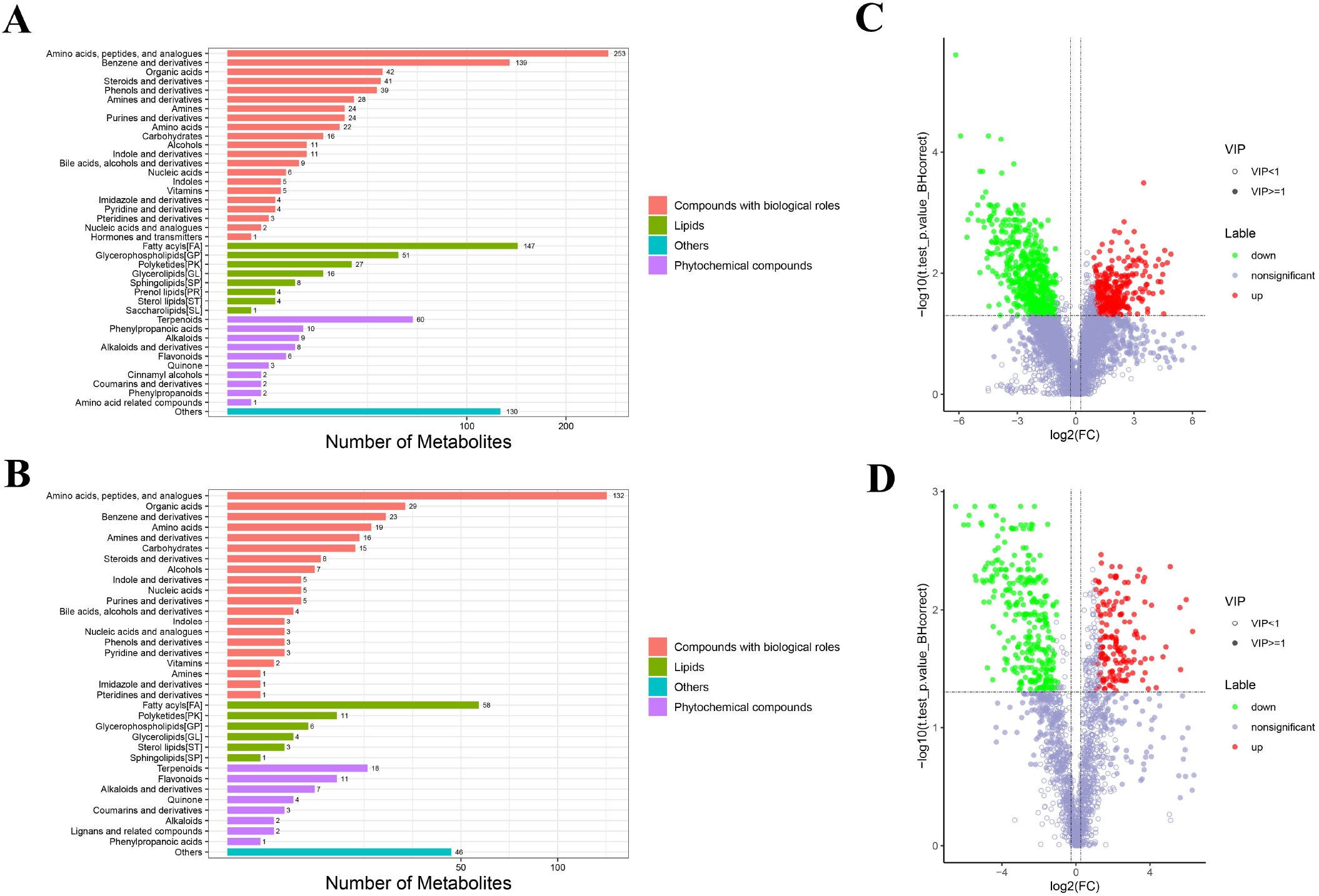
Taxonomic annotation of identified metabolites from the intestine of *M. japonicus*. **(A)** positive ion mode. **(B)** negative ion mode. Others means that the classification information is other categories, and the identified metabolites without classification information do not participate in the statistics.

#### 3.4.2. Identification and Functional Annotation of the DMs

The DMs between the control group and DIV1-infected group were identified by the PLS-DA model with a cut-off VIP ≥ 1 and *q*-value ≤ 0.05. In the positive ion mode, a total of 868 DMs were obtained, including 312 up-regulated DMs and 556 down-regulated DMs, of which 419 DMs were identified in the database (Figure 6C). In the negative ion mode, a total of 454 DMs were obtained, including 158 up-regulated DMs and 296 down-regulated DMs, of which 209 DMs were identified in the database (Figure 6D).

For the KEGG pathway enrichment analysis, the top 20 KEGG pathway enrichments influenced by DIV1 infection in the positive ion mode and negative ion mode were shown in Figure 7A and B, respectively. Among of them, four KEGG pathways related to vitamin metabolism were significantly enriched, including “vitamin digestion and absorption”, “retinol metabolism”, “vitamin B6 metabolism” and “pantothenate and CoA biosynthesis”. Three KEGG pathways related to amino acid metabolism were significantly enriched, including “tyrosine metabolism”, “phenylalanine metabolism” and “alanine, aspartate and glutamate metabolism”. Two Warburg effect marker pathways were significantly enriched, including “pyruvate metabolism” and “glycolysis / gluconeogenesis”. Both in positive ion mode and in negative ion mode, “primary bile acid biosynthesis”, “Bile secretion”, “metabolic pathways”, “Linoleic acid metabolism” and “arachidonic acid metabolism” were significantly enriched in KEGG pathway enrichment analysis.

**Figure 7.**
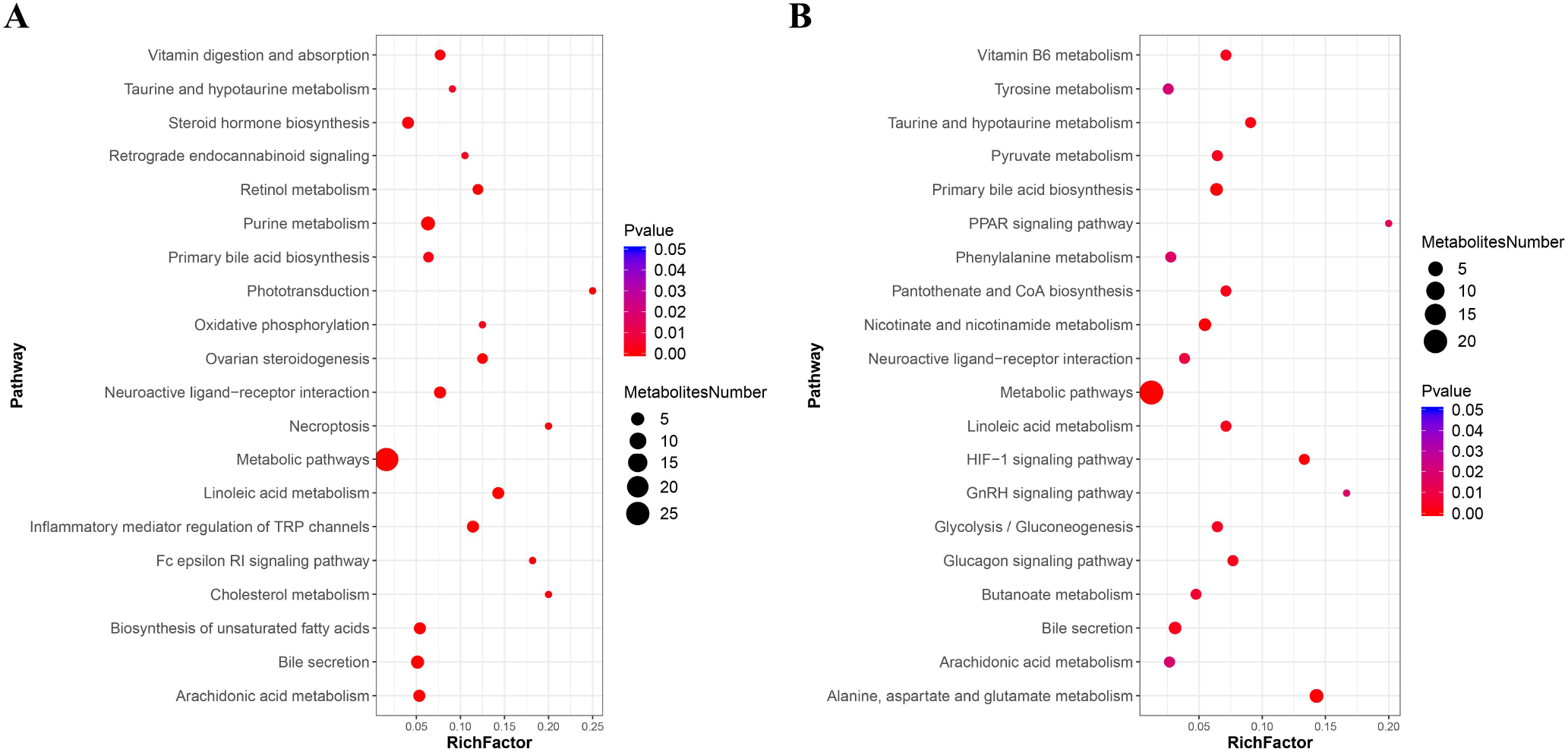
KEGG pathway enrichment analysis of DMs between the control group and DIV1-infected group. **(A)** positive ion mode. **(B)** negative ion mode.

Further, in the positive ion mode, total of 41 known DMs were significantly enriched, including 25 up-regulated DMs and 16 down-regulated DMs (Table 2); in the negative ion mode, a total 24 DMs were significantly enriched, including 17 up-regulated DMs and 7 down-regulated DMs (Table 3). It was worth noting that seven types of lipid fatty acids were significantly changed after DIV1 infection, including arachidonic acid, eicosapentaenoic acid, docosapentaenoic acid, 9-oxo-10(e),12(e)-octadecadienoic acid, gamma-linolenic acid, 8(s)-hydroxy-(5z,9e,11z,14z) eicosatetraenoic acid and 16-hydroxyhexadecanoic acid. In addition, two Warburg effect marker metabolites were significantly up-regulated, including L-(+)-lactic acid and Pyruvic acid.

**Table 2.**
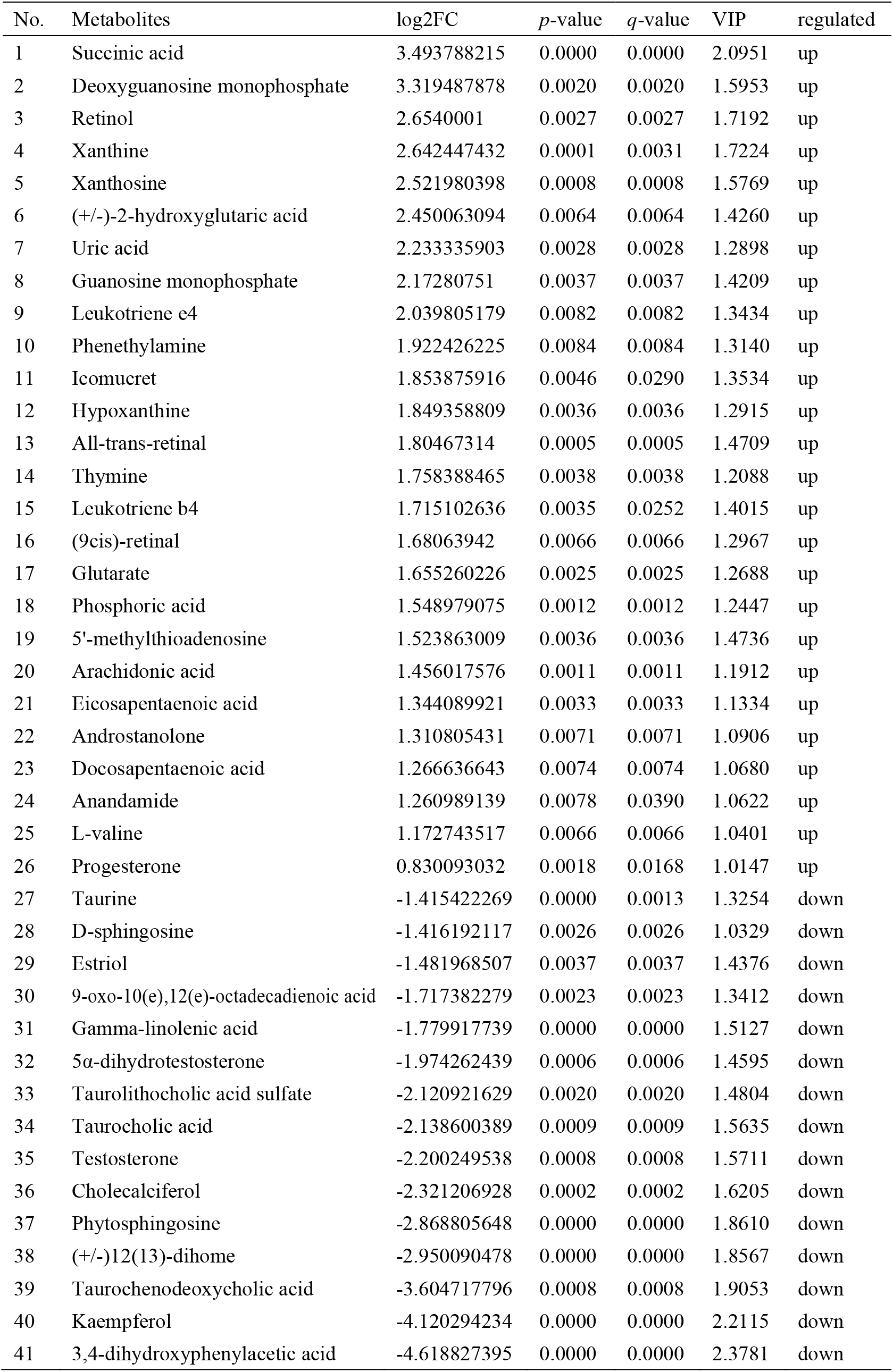
41 known differential metabolites were significantly enriched in KEGG pathway enrichment analysis in positive ion mode.

**Table 3.** 24 known differential metabolites were significantly enriched in KEGG pathway enrichment analysis in negative ion mode.

| No. | Metabolites | log2FC | *p*-value | *q*-value | VIP | regulated |
| --- | --- | --- | --- | --- | --- | --- |
| 1 | Cytidine | 3.146492307 | 0.0011 | 0.0109 | 1.7973 | up |
| 2 | Pantothenic acid | 2.722553458 | 0.0069 | 0.0314 | 1.3379 | up |
| 3 | Pyridoxine | 2.709555326 | 0.0098 | 0.0394 | 1.3698 | up |
| 4 | N-acetyl-l-phenylalanine | 2.593234239 | 0.0055 | 0.0275 | 1.4078 | up |
| 5 | Argininosuccinic acid | 2.556306784 | 0.0053 | 0.0271 | 1.3755 | up |
| 6 | Succinic semialdehyde | 2.279560440 | 0.0037 | 0.0224 | 1.3252 | up |
| 7 | Uric acid | 2.258820397 | 0.0005 | 0.0072 | 1.4234 | up |
| 8 | 8(s)-hydroxy-(5z,9e,11z,14z)-eicosatetraenoic acid | 2.183581791 | 0.0027 | 0.0191 | 1.3515 | up |
| 9 | Phosphoric acid | 2.161629747 | 0.0006 | 0.0085 | 1.4491 | up |
| 10 | Adenine | 1.969196548 | 0.0076 | 0.0333 | 1.1510 | up |
| 11 | L-(+)-lactic acid | 1.909043413 | 0.0023 | 0.0172 | 1.2639 | up |
| 12 | Niacin | 1.536649754 | 0.0141 | 0.0473 | 1.1171 | up |
| 13 | Pyruvic acid | 1.437494091 | 0.0040 | 0.0236 | 1.0153 | up |
| 14 | Terephthalic acid | 1.360645202 | 0.0001 | 0.0034 | 1.0953 | up |
| 15 | Taurine | 1.331303934 | 0.0080 | 0.0344 | 1.0127 | up |
| 16 | Arachidonic acid | 1.306845500 | 0.0025 | 0.0182 | 1.0367 | up |
| 17 | N-acetyl-l-aspartylglutamic acid | 1.085968454 | 0.0004 | 0.0067 | 1.0802 | up |
| 18 | N-acetyl-d-mannosamine | -1.046027092 | 0.0011 | 0.0110 | 1.0350 | down |
| 19 | 16-hydroxyhexadecanoic acid | -1.569179503 | 0.0015 | 0.0129 | 1.3084 | down |
| 20 | Cholic acid | -2.838563734 | 0.0000 | 0.0021 | 1.6745 | down |
| 21 | (+/-)12(13)-dihome | -2.975964065 | 0.0000 | 0.0019 | 1.7010 | down |
| 22 | Luteolin | -3.930160375 | 0.0000 | 0.0017 | 1.9612 | down |
| 23 | Taurochenodeoxycholic acid | -4.316168826 | 0.0001 | 0.0038 | 2.1122 | down |
| 24 | N-acetylmuramic acid | -4.952322025 | 0.0001 | 0.0046 | 1.9742 | down |

### 3.5. Association Between the Altered Metabolites and Microbial

To reveal the relationships between intestinal microbial and DMs, scatterplots at the phylum, family, and genus levels were generated by canonical correlation analysis (Figure S2). The results showed that there was a strong correlation (R value > 0.75) between the host intestinal microbial and DMs at all taxonomic levels, and the samples from different groups had a large degree of dispersion. To further analyze the relationship between the nine marker DMs (including seven types of fatty acids and two Warburg effect marker metabolites) and host intestinal microbial, heat maps at the phylum, family, and genus levels were generated by spearman correlation analysis (Figure 8). The results showed that, at the phylum classification level, Proteobacteria was positively associated with arachidonic acid, 8(s)-hydroxy-(5z,9e,11z,14z) eicosatetraenoic acid, docosapentaenoic acid, eicosapentaenoic acid, pyruvic acid and L-(+)-lactic acid, and negatively associated with gamma-linolenic acid, 16-hydroxyhexadecanoic acid and 9-oxo-10(e),12(e)-octadecadienoic acid. The association of Cyanobacteria and Actinobacteria with marker metabolites was completely opposite to that of Proteobacteria, while neither Bacteriodetes nor Firmicutes were significantly associated with the marker metabolites (*p* > 0.05). It was worth noting that, *Photobacterium* and *Vibrio* in the Vibrionaceae that dominated the DIV1-infected group, were positively associated with Warburg effect marker metabolites L-(+)-lactic acid and pyruvic acid. In contrast, *Corynebacterium* in the Corynebacteriaceae that dominated the control group, was negatively correlated with L-(+)-lactic acid and pyruvic acid. This phenomenon also appeared in the correlation with fatty acid metabolites.

**Figure 8.**
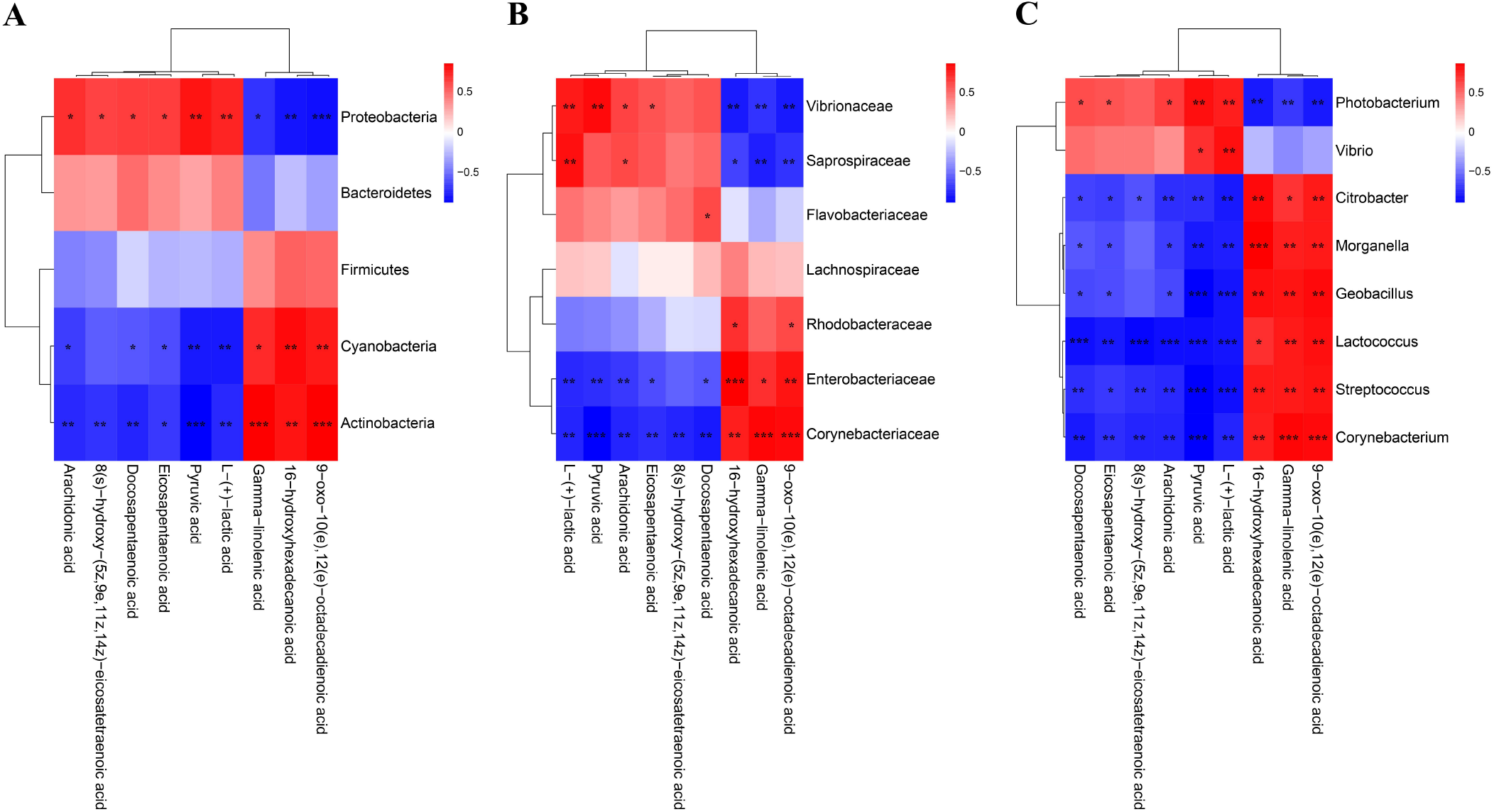
Heap maps of correlation between the nine marker DMs and the hosts intestinal microbial. **(A)** Phylum classification level. **(B)** Family classification level. **(C)** Genus classification level. Red indicateds a positive coreelation, and blue represents a negative correlation. * Indicates a significant difference (\**p* < 0.05; \*\**p* < 0.01; \*\*\**p* < 0.001).

## 4. Discussion

Growing evidences suggested that viral infection or environmental stress can lead to changes in the structure and function of the host’s intestinal microbiota, and further affect the normal metabolism of the host and caused various adverse reactions (Hsu et al., 2019; Qiao et al., 2019; Duan et al., 2021a; Duan et al., 2021b). To date, there are no reports on the effects of DIV1 infection on the intestinal microbiota and metabolites of *M. japonicus*. The present study provides insights into the interaction of *M. japonicus* and DIV1 through the histological analysis, enzyme activity analysis and the integrated analysis of intestinal microbiome and metabolomics.

The results of histological analysis showed that severe intestinal mucosal damage was observed in the intestine of DIV1-infected shrimp, and the intestinal epithelial cells were completely detached from the basement membrane. Similarly, Xue et al. found that WSSV-infected crayfish *Procambarus clarkii* exhibited worse intestinal histomorphology, with thinner intestinal walls and shorter intestinal villi compared to healthy crayfish (Xue et al., 2022). Liang et al. found that the intestinal epithelial cells of *L. vannamei* infected with *Vibrio parahaemolyticus* were completely detached from the basement membrane (Liang et al., 2020). Except for the intestinal mucosal damage, the disturbance of intestinal immunity and digestive function was also observed in the DIV1-infected shrimp. Enzyme activity analysis showed that all all digestive enzymes and immune enzymes were significantly altered (*p* < 0.001), indicating that DIV1 infection severely affected the normal immune and digestive functions of the *M. japonicus* intestine. Studies have found that shrimp hemocytes can produce a large amount of free radicals and reactive oxygen species (ROS) during phagocytosis, thereby effectively killing the invading pathogens (Bogdan et al., 2000; Duan et al., 2015). However, the mass accumulation of ROS in animals will cause serious cell damage, resulting in various diseases (Yu et al., 1994). SOD and CAT are two kinds of antioxidant enzymes, which can protect cells against <u>oxidative stress</u> (Zhu et al., 2012; Wu et al., 2014; Chen et al., 2015). In our study, the activities of SOD was significantly increased in the intestine of the *M. japonicus* under DIV1 infection, while the activities of CAT was significantly decreased. It meant that DIV1 infection may disrupt the oxidative and antioxidant balance in the shrimp intestine. LYZ has been shown to play an important role in shrimp resistance to viral and bacterial infections (Kaizu et al., 2011; Karthik et al., 2014; Liu et al., 2016). The activity of LYZ can directly reflect the strength of shrimp immune function. After infection with DIV1, the acitivity of LYZ in the intestine of *M. japonicus* was significantly decreased, which indicated that the intestinal immune function was suppressed to some extent. α-AMS, LPS and TPS are three important digestive enzymes. After infection with DIV1, the activities of intestinal digestive enzymes showed different trends, suggesting that DIV1 infection caused the digestive dysfunction of shrimp. This may be one of the reasons for the decreased feeding rate of DIV1-infected shrimp and caused the symptoms of empty stomach and intestine (Chen et al., 2019; Qiu et al., 2019; Liao et al., 2020; He et al., 2021a; He et al., 2021b).

In intestinal microbiom analysis, after DIV1 infection at 24 h, the intestine microbiome was significantly changed in the composition and diversity at the phylum, family and genus levels. In this study, the intestinal microbiota in *M. japonicus* were dominated by five phyla of Proteobacteria, Actinobacteria, Bacteroidetes, Firmicutes and Cyanobacteria in both the control groups and DIV1-infected group, which is consistent with the previous microbiome studies in *L. vannamei*. At the family level, the relative abundance of Vibrionaceae significantly increased in the intestine of DIV1-infected *M. japonicus* compared with healthy *M. japonicus*, which mainly reflected in the significant increase in the relative abundance of *Photobacterium* and *Vibrio* at the genus level. In addition, both the LEfSe cladogram and LDA score of LEfSe-PICRUSt showed that *Photobacterium* and *Vibrio* dominated in the DIV1-infected group. Several previous studies have shown that *Photobacterium* can cause various diseases in fish and shrimp (Vaseeharan et al., 2007; Kanchanopas-Barnette et al., 2009; Zhang et al., 2011; Singaravel et al., 2020; Wang et al., 2020), causing huge losses in aquaculture. *Vibrio harveyi, Vibrio alginolyticus* and *V. parahaemolyticus* are three species of *Vibrio*, which are considered to be the main opportunistic pathogens in shrimp disease outbreaks (Zhou et al., 2012; Santhyia et al., 2015; Nguyen et al.,2021). These results suggested that intestinal pathogenic bacteria were increased after DIV1 infection, which may probably because the DIV1 infection disrupted the mechanical barrier of the intestine and impaired the ability of intestine to select microorganisms. *Photobacterium* and *Vibrio* may serve as marker microbial taxa for DIV1 infection. The results of PICRUSt functional prediction showed that the relative abundance of “Carbohydrate metabolism”, “Metabolism of cofactors and vitamins” and “Amino acid metabolism” were the top 3 in the two groups. It meant that intestinal microbiota played an important role in regulating host vitamin metabolism, carbohydrate metabolism and amino acid metabolism. Notably, the relative abundance of “infectious diseases: bacterial” was significantly increased under DIV1 infection, which further suggested that DIV1 infection could cause secondary bacterial infection.

A recent study found that the “Linoleic acid metabolism” and “arachidonic acid metabolism” were also the most disturbed pathways by HCoV-299E infection (Yan et al., 2019). Interestingly, exogenous supplement of linoleic acid or arachidonic acid in HCoV-229E-infected cells significantly suppressed HCoV-229E virus replication (Yan et al., 2019). Another study showed that linoleic acid could directly bound to WSSV to inhibited the viral replication and indirectly participate in the immune response against WSSV by activating the ERK-NF-κB signaling pathway to promote the expression of antimicrobial peptides and IFN-like gene Vago5 (Li et al., 2021). Consistently, the intestinal metabolomics analysis in this study showed that lipid metabolism in the intestine of *M. japonicus* was remodeled after DIV1 infection, and a total of seven types of lipid fatty acids were significantly changed, including arachidonic acid, eicosapentaenoic acid, docosapentaenoic acid, 9-oxo-10(e), 12(e) -octadecadienoic acid, gamma-linolenic acid, 8(s)-hydroxy-(5z,9e,11z,14z) eicosatetraenoic acid and 16-hydroxyhexadecanoic acid. In addition, two Warburg effect marker metabolites were significantly up-regulated, including L-(+)-lactic acid and Pyruvic acid. KEGG enrichment analysis indicated that the “Linoleic acid metabolism” and “arachidonic acid metabolism” pathways were the most affected by DIV1 infection. These results suggested that the lipid metabolic reprogramming of *M. japonicus* was significantly associated with DIV1 infection and replication. After infected with DIV1, two marker metabolites of the Warburg effect were significantly up-regulated in the shrimp intestine, including L-(+)-lactic acid and Pyruvic acid, and significantly enriched in the “pyruvate metabolism” and “glycolysis / gluconeogenesis” pathways. In addition, some amino acid metabolism-related pathways were also significantly enriched under DIV1 infection, including “tyrosine metabolism”, “phenylalanine metabolism” and “alanine, aspartate and glutamate metabolism”. These phenomena also appeared in previous studies (Liao et al., 2020; He et al., 2021b; He et al., 2022), which may be attributed to the material and energy requirements of DIV1 replication and suggested that DIV1 may contain AMGs for regulating host amino acid metabolism, carbohydrate metabolism and energy metabolism. It was worth noting that some pathways related to vitamin metabolism were significantly enriched, such as “retinol metabolism”, “vitamin B6 metabolism” and “pantothenate and CoA biosynthesis”. Retinol, also known as vitamin A (VA), plays an important and positive role in intestinal mucosal immunity and repair (Sirisinha., 2015). The B vitamins can regulate the energy metabolism, nutrient accumulation and immune defense function of aquatic animals (Mai et al., 2000). Our previous study showed that DIV1 infection can regulate the vitamin metabolism of shrimp by affecting the expression of intestinal miRNA and mRNA (Liao et al., 2020; He et al., 2022). In this study, from the metabolome level, it was further demonstrated that DIV1 infection can regulate vitamin metabolism in the intestine, thereby affecting the normal physiological function of *M. japonicus*.

Intestinal microbiota variation caused by viral infection or environmental stress can affect the metabolism and immunity of their host, and increase disease susceptibility (Levy et al., 2017; Holt et al., 2020). To futher expore the relationship between intestine microbiota variation and metabolites, the correlation analysis of dominant intestinal microbial and marker DMs was carried out. In this study, the increased levels of *Photobacterium* and *Vibrio* were positively correlated with changes in Warburg effect marker metabolites L-(+)-lactic acid and pyruvic acid. This indicated that *Photobacterium* and *Vibrio* synergize with DIV1 to jointly promoted the Warburg effect, providing material and energy for the successful replication of DIV1. In addition, the abundances of *Photobacterium* and *Vibrio* were positively associated with arachidonic acid, eicosapentaenoic acid, docosapentaenoic acid and 8(s)-hydroxy-(5z,9e,11z,14z) eicosatetraenoic acid and negatively associated with 9-oxo-10(e),12(e)-octadecadienoic acid, gamma-linolenic acid and 16-hydroxyhexadecanoic acid. However, *Corynebacterium*, which dominated the control group, was just the opposite of the above. It suggested that *Photobacterium* and *Vibrio* were involved in the metabolic reprogramming of the shrimp intestine after DIV1 infection. Several of the highly relevant bacteria and metabolites selected in this study might be used as biomarkers for shrimp in response to DIV1 infection.

## 5. Conclusions

In conclusion, through the histological analysis, enzyme activity analysis and the integrated analysis of intestinal microbiome and metabolomics, the present study revealed that DIV1 infection can lead to the damage of the intestinal mechanical barrier of shrimp, the imbalance of oxidative and antioxidant capacity, the decrease of immune enzyme activities, and the disturbance of digestive enzyme activities, thereby causing secondary bacterial infections, including *Photobacterium* and *Vibrio*. At the same time, harmful bacteria can cooperate with DIV1 to promote the Warburg effect and induce metabolic reprogramming, thereby creating favorable conditions for the replication of DIV1 (Figure 9). This study is the first to report the changes of intestinal microbiota and metabolites of *M. japonicus* under DIV1 infection, demonstrating that DIV1 can induce secondary bacterial infection and metabolic reprogramming, and several highly related bacteria and metabolites were screened as biomarkers. These biomarkers can be leveraged for diagnosis of pathogenic infections or incorporated as exogenous metabolites to enhance immune response.

**Figure 9.**
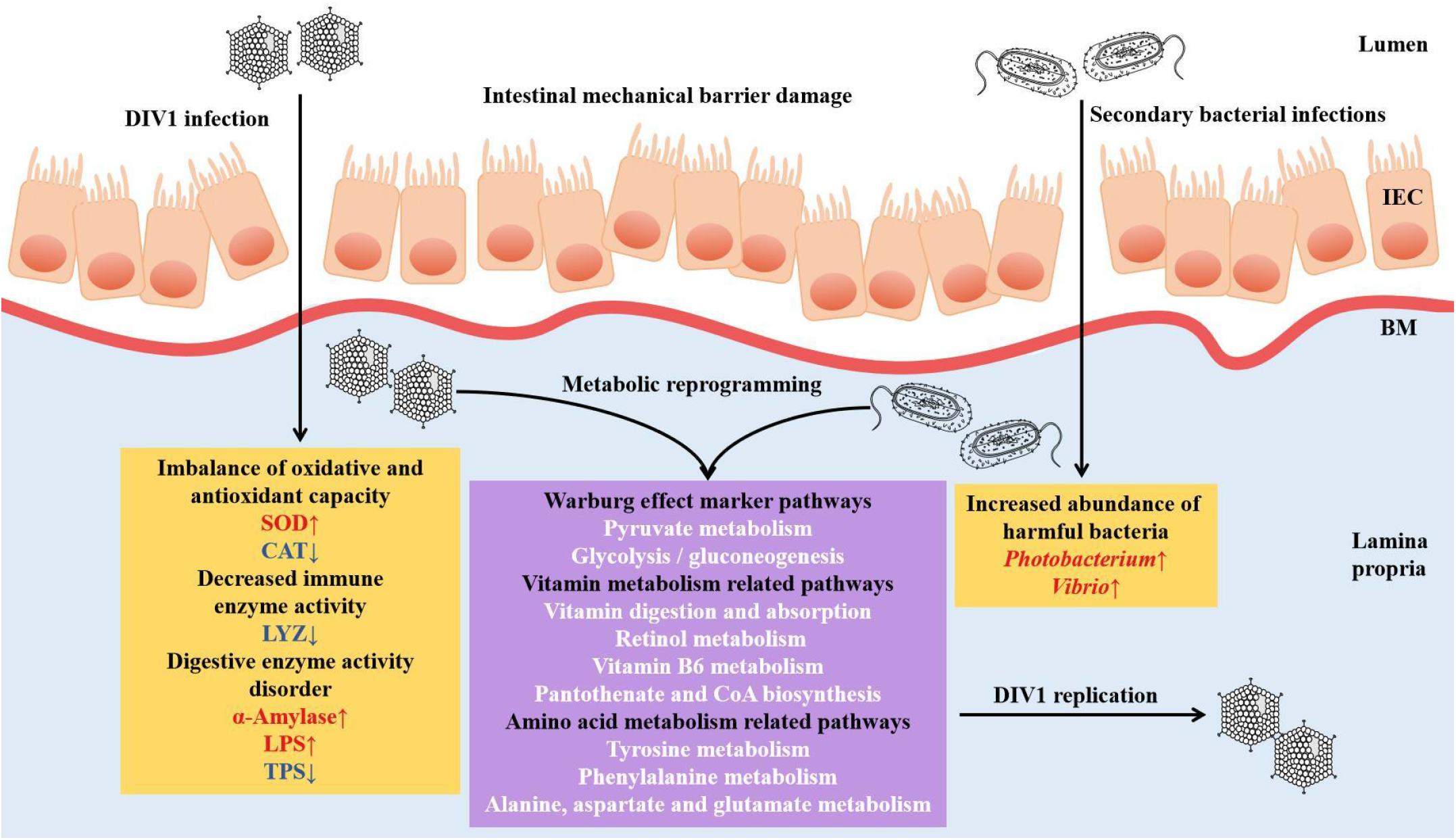
Proposed schematic diagram of secondary bacterial infection and metabolic reprogramming of *M. japonicus* induced by DIV1 infection. IEC: intestinal epithelial cells; BM: basement membrane; SOD: superoxide dismutase; CAT: catalase; LYZ: lysozyme; LPS: lipase; TPS: trypsin.

## Acknowledgments

Our acknowledgments go to all funders of this work.

## Figure Legends

**Figure S1. Derived PLS-DA score plots and corresponding permutation testing of PLS-DA from the LC-MS metabolite profiles in the intestine of *M. japonicus* after DIV1 infection. (A)** PLS-DA score plot of positive ion mode. **(B)** PLS-DA score plot of negative ion mode. **(C)** Permutation testing of positive ion mode. **(D)** Permutation testing of negative ion mode.

**Figure S2. Scatter plots of correlation between DMs and hosts intestinal microbial. (A)** Phylum classification level. **(B)** Family classification level. **(C)** Genus classification level.

## Highlights

- DIV1 infection disrupts the shrimp intestinal mechanical barrier.
- DIV1 infection resulted in imbalance of oxidative and antioxidant capacity, decreased immune enzyme activity, and disordered digestive enzyme activity in shrimp.
- DIV1 infection caused secondary bacterial infections, including *Photobacterium* and *Vibrio*.
- DIV1 can cooperate with harmful bacteria to induce metabolic reprogramming.

